# Tissue-specific chromatin binding patterns of *C. elegans* heterochromatin proteins HPL-1 and HPL-2 reveal differential roles in the regulation of gene expression

**DOI:** 10.1101/2023.01.13.523961

**Authors:** Patricia de la Cruz Ruiz, María Jesús Rodríguez-Palero, Peter Askjaer, Marta Artal-Sanz

## Abstract

Heterochromatin is characterized by an enrichment of repetitive elements and low gene density and is often maintained in a repressed state across cell division and differentiation. The silencing is mainly regulated by repressive histone marks, such as H3K9 and H3K27 methylated forms and the heterochromatin protein 1 (HP1) family. Here, we analyzed in a tissue-specific manner the binding profile of the two HP1 homologs in *Caenorhabditis elegans*, HPL-1 and HPL-2, at the L4 developmental stage. We identified the genome-wide binding profile of intestinal and hypodermal HPL-2 and intestinal HPL-1 and compared them to heterochromatin marks and other features. HPL-2 associated preferentially to the distal arms of autosomes and correlated positively with methylated forms of H3K9 and H3K27. HPL-1 was also enriched in regions containing H3K9me3 and H3K27me3 but exhibited a more even distribution between autosome arms and centers. HPL-2 showed a differential tissue-specific enrichment for repetitive elements, conversely with HPL-1 that exhibited a poor association. Finally, we found a significant intersection of genomic regions bound by the BLMP-1/PRDM1 transcription factor and intestinal HPL-1, suggesting a co-repression role during cell differentiation. Our study uncovers both shared and singular properties of conserved HP1 proteins, providing information about genomic binding preferences in relation to their role as heterochromatic markers.

## Introduction

Heterochromatin in worms, like in other species, is tightly packaged, presents a low gene density and is usually enriched at the nuclear periphery (Guand Fire 2010; Ahringer and Gasser 2018). Facultative heterochromatin is defined by the presence of histone H3 lysine 27 tri-methylation (H3K27me3), which can be erased by demethylation during development leading to a transcriptionally active state. The Polycomb PRC2 complex comprised by MES-2, MES-3, and MES-6 proteins is responsible for H3K27 methylation (Steffen and Ringrose 2014; Piunti and Shilatifard 2016). The constitutive form of heterochromatin is characterized by H3K9me3, which ensures homeostasis and differentiation of tissues across species (Padeken *et al*. 2022). H3K9 methylation is performed in sequential steps by MET-2/SETDB1 enzyme that carries out mono- and di-methylation and SET-25/G9a, in charge of tri-methylation. Both enzymes are main players in the perinuclear attachment of H3K9me enriched heterochromatin (Towbin *et al*. 2012). These heterochromatin domains usually reside within the chromosomal distal regions (“arms”), where most meiotic recombination occurs. The arms are also enriched in repetitive DNA elements, decorated by H3K9me2 and/or H3K9me3 (Consortium 1998; Liu *et al*. 2011; McMurchy *et al*. 2017). On the contrary, genes in autosome centers are generally higher expressed and more evolutionarily conserved (Consortium 1998).

In *C. elegans*, HPL-2 is considered as a marker for heterochromatin domains in developing embryos (Grant *et al*. 2010). HPL-2 is part of the heterochromatin protein 1 (HP1) family, which is highly conserved across the eukaryotic kingdom. HP1 was first characterized in fruit flies, as a component responsible for variegation position effect, in which inhibition of gene expression occurs via imposition of heterochromatin-like structure to inactivate it (Eissenberg *et al*. 1990). The two HP1 homologues in *C. elegans*, HPL-1 and HPL-2, possess a conserved structure that consists of an N-terminal chromo domain (CD) involved in binding to the K9 residue on histone H3 in its mono, di or tri-methylated form. The CD is followed by a flexible hinge region and a C-terminal chromo shadow domain (CSD) required for dimerization and interaction with other proteins (Eissenberg 2001; Schott *et al*. 2006).

HPL-1 and HPL-2 are 48% identical along their entire length, with the highest homology within the CD and CSD (60%; Figure S1C-D). The hinge region is more divergent and consists of only 19 amino acid residues in HPL-1 versus 31 residues in HPL-2. Their expression is ubiquitous in nuclei of most cells throughout development (Couteau *et al*. 2002; Schott *et al*. 2006). The observation of post-embryonic defects in *hpl-2* mutants, such as sterility, defective growth and brood size, larval arrest and multivulva phenotypes demonstrated that HPL-2 functions in cell fate determination and gonad development. The similar synthetic phenotypes presented in double *hpl-1;hpl-2* mutants prompted to conclude a redundant functionality for HPL-1, which did not show obvious phenotypes when mutated alone (Cardoso *et al*. 2005; Coustham *et al*. 2006; Schott *et al*. 2006; Simonet *et al*. 2007; Schott *et al*. 2009; Koester-Eiserfunke and Fischle 2011). HPL-2 has also been implicated in DNA replication and stress response, being enriched in DNA repetitive elements (Cardoso *et al*. 2005; Coustham *et al*. 2006; Schott *et al*. 2009; Black *et al*. 2010; Meister *et al*. 2011; Garrigues *et al*. 2015).

Comparatively, little is known about HPL-1 functions in *C. elegans*. A physical interaction between HPL-1 and the mono-methylated H1K14me1 histone variant has been observed to regulate innate immunity against pathogenic bacterial infection (Studencka *et al*. 2012a). Like HPL-2, loss of HPL-1 results in transcriptional alteration of genes that encode nuclear hormone receptor family proteins such as NHR-60, NHR-156, transcription factors including MIZ-1, ZIP-3, KZIP-8, MADF-2, homeobox genes like CEH-82 or the homeodomain gene LIM-7, that cause male tail defects (Studencka *et al*. 2012b). Recently, HPL-1 was related to transgenerational embryonic lethality under specific paternal DNA damage. HPL-1 depletion induced a reduction in H3K9me2 levels in the germline, which was beneficial for F2 generation survival (Wang *et al*. 2022).

Despite the association of HP1 proteins with different functions, tissue-specific behavior of HPL-1 and HPL-2 has not been characterized. These types of studies have become important to investigate heterogeneity between tissues. This may help to understand, for instance, potential regulatory links between epigenetic alterations and target gene expression changes to establish an integrative analysis (Yilmaz *et al*. 2020). To reinforce this idea, contrasting results have been reported when performing rescue experiments with HP1 proteins upon endoplasmic reticulum stress. For example, upon whole-body loss of HPL-2, intestinal-specific expression of HPL-2 restored wild type level of the stress response, while it was detrimental upon neuronal-specific expression (Kozlowski *et al*. 2014). However, overexpression in either of the two tissues, was sufficient to provide partial rescue of dauer exit (Meister *et al*. 2011). This highlights differences in tissue response to lack of HPL-2 when evaluating distinct pathways.

To contribute to the characterization of HP1 proteins, a comparative study of HPL-1 and HPL-2 homologous proteins in *C. elegans* in a tissue-specific manner is reported here. We identified specific genomic regions bound by HPL-2 in hypodermal and intestinal tissues, as well as HPL-1-associated regions in the intestine at the L4 developmental stage. We found that HPL-2 associated preferentially to chromosome distal arms, while HPL-1 was more equally distributed between autosome arms and center. Moreover, both HP1 homologs strongly correlated with the two repressive histone marks H3K9me3 and H3K27me3 in both tissues analyzed. However, only HPL-2 binding pattern coincided with mono- and di-methylated H3K9 forms.

We confirmed a general repressive role for HP1 proteins when comparing with transcribed genes in the tissues studied. In addition, the large number of unique genes, together with gene ontology analyses, suggest tissue-specific patterns of gene silencing. Moreover, we discovered a potential co-regulation of chromatin regions by BLMP-1/PRDM1 transcription factor and intestinal HPL-1. Lastly, HPL-2 was found to be enriched at DNA repetitive elements in both tissues, conversely to intestinal HPL-1. In summary, our study uncovered both unique and common features of HP1 *C. elegans* homologous proteins in two differentiated tissues.

## Results

### HPL-1 and HPL-2 differentially associate to the genome

To study the function of *C. elegans* HP1 proteins, the DNA binding profiles for HPL-1 and HPL-2 were determined by tissue-specific DNA adenine methylation identification (DamID) at the L4 larval stage {de la Cruz Ruiz, 2022 #106}. We generated Dam::HPL-1/2 fusions by inserting Dam at the sites previously used for multicopy (Couteau *et al*. 2002) and endogenous (Patel and Hobert 2017) tagging of HPL-1/2, which results in functional nuclear-localised proteins (Figure S1A). Of the three *hpl-2* isoforms, our Dam insertion tags specifically *hpl-2a*, the most expressed isoform based on RNAseq data (Figure S1B-C). Ubiquitous expression of fluorescent protein fusions at endogenous levels results in phenotypically wild type animals whereas Dam fusions are expressed at much lower levels from a non-induced *hsp-16*.*41* promoter and only in either intestinal or hypodermal cells. We focused on the intestine, a major metabolic tissue of *C. elegans* amenable to this technique (Cabianca *et al*. 2019), and included also the hypodermis for HPL-2 to compare binding profiles across two tissues. We confirmed the tissue-specific expression of intestinal Dam::HPL-1 and Dam::HPL-2 and hypodermal Dam::HPL-2 (Figure S2), as well as the correlation between replicas (Figure S3). We then first analyzed the results at 100 kb resolution to obtain a broad overview of the behavior of the two HP1 proteins. In agreement with previous observations in embryonic cells, HPL-2 is enriched in distal regions (“arms”) of autosomes in both hypodermal and intestinal tissues and depleted from autosome centers (Figure 1A; Figure S4A)(Garrigues *et al*. 2015). We found that chromosome arms with meiotic pairing center were more frequently in contact with HPL-2 in the hypodermis compared to the intestine (Figure 1A). Surprisingly, HPL-1 in the intestine is distributed more equally between autosome arms and centers, being significantly different from HPL-2 (Figure 1A). The difference between HPL-1 and HPL-2 binding profiles in the intestine is also observed at higher resolution (10 kb) for all autosomes (Figure 1B; Figure S4A). This suggests that HPL-1 might have unique roles in *C. elegans*. In addition, both HPL-2 DamID profiles exhibit a significant overlap with previous whole-animal HPL-2 ChIP datasets (Gerstein *et al*. 2010)(Figure S5 A-C; Table S1).

**Figure 1.**
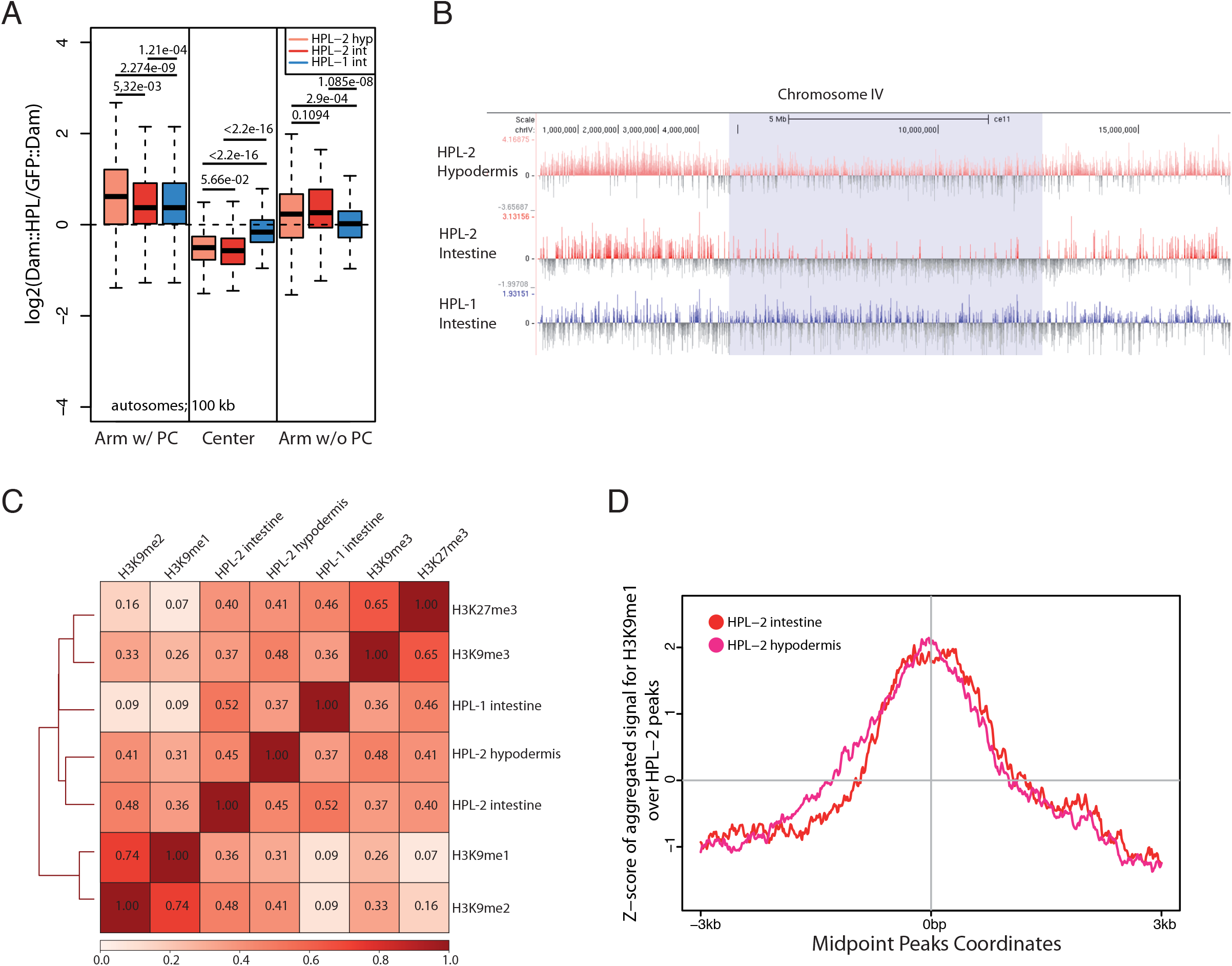
Tissue-specific chromatin binding of HPL-1 and HPL-2 and their association to histone marks. (A) Boxplot showing HP1 association profile in all autosomes based on arms with and without meiotic pairing centers at 100 kb bin resolution in hypodermal and intestinal tissues. p-values from Wilcoxon rank sum test with continuity correction are indicated. PC: pairing center. (B) Example of average signal tracks over 10 kb regions in chromosome IV for HP1 proteins in hypodermal and intestinal tissues. Light blue region represents chromosome center, while flanking regions correspond to left and right arms. Snapshot from IGV viewer. (C) Heatmap showing pairwise Pearson correlation coefficients between indicated datasets at 25 bp window size. (D) Aggregation plot depicting the genome-wide average signal tracks for H3K9me1 dataset over HPL-2 hypodermal and intestinal peaks. Center position represents the midpoint location of peaks. Regions up to 3 kb upstream and 3 kb downstream of the midpoint in 10 bp bins are shown.

### HPL-1 and HPL-2 differentially correlate with H3K9me3 and H3K27me3 histone marks

As reader of methylated H3K9, HPL-2 binding sites in the genome coincide with the pattern of mono and di-methylated H3K9 (H3K9me1-2) in *C. elegans* embryos, and less so with H3K9me3 (Garrigues *et al*. 2015). To determine if the correlation also exists in differentiated cells and for both HP1 proteins, we compared the HPL-1 and HPL-2 binding profiles with published histone ChIP datasets from the L3 larval stage. Indeed, a high correlation was found between both HP1 proteins and repressive H3K9me3 and H3K27me3 histone marks, clustering together in the pairwise Pearson correlation heatmap (Figure 1C). Both histone marks usually colocalize on distal autosome arms, while only H3K27me3 associates with central regions as well (Liu *et al*. 2011; Ahringer and Gasser 2018).

Interestingly, HPL-2 in both tissues positively correlated with mono and di-methylated H3K9, whereas HPL-1 protein failed to show any significant association (Figure 1 C-D, Figure S4B-D). Therefore, our data indicates that only HPL-2 exhibits more flexibility to bind to the three methylated forms of H3 in both differentiated tissues, suggesting a differential role in the control of gene expression for HPL-1 and HPL-2 through development.

### HP1 proteins bind to genomic regions in a tissue-specific manner

We next asked if the differences in distribution of large HP1-associated genomic regions across autosomes (Figure 1A-B) and the correlation with heterochromatin marks (Figure 1C-D) reflect a differential binding of HPL-1 and HPL-2 to smaller specific DNA regions. In addition, the different proportion of overlapping peaks between HP1 proteins and whole-animal HPL-2 ChIP data could indicate tissue-specific differences (Figure S5A; Table S1). To address this, we compared the regions present in each HP1 dataset. Strikingly, we found that around 70% of bound genomic regions are unique for either HPL-1 or HPL-2 (Figure 2A). The highest overlap was found between the two tissue-specific HPL-2 profiles, suggesting a conserved regulation for specific regions in both tissues (Figure 2A). However, the proportion of overlapped regions between all datasets does not reach 30%, remarking the high degree of specificity.

**Figure 2.**
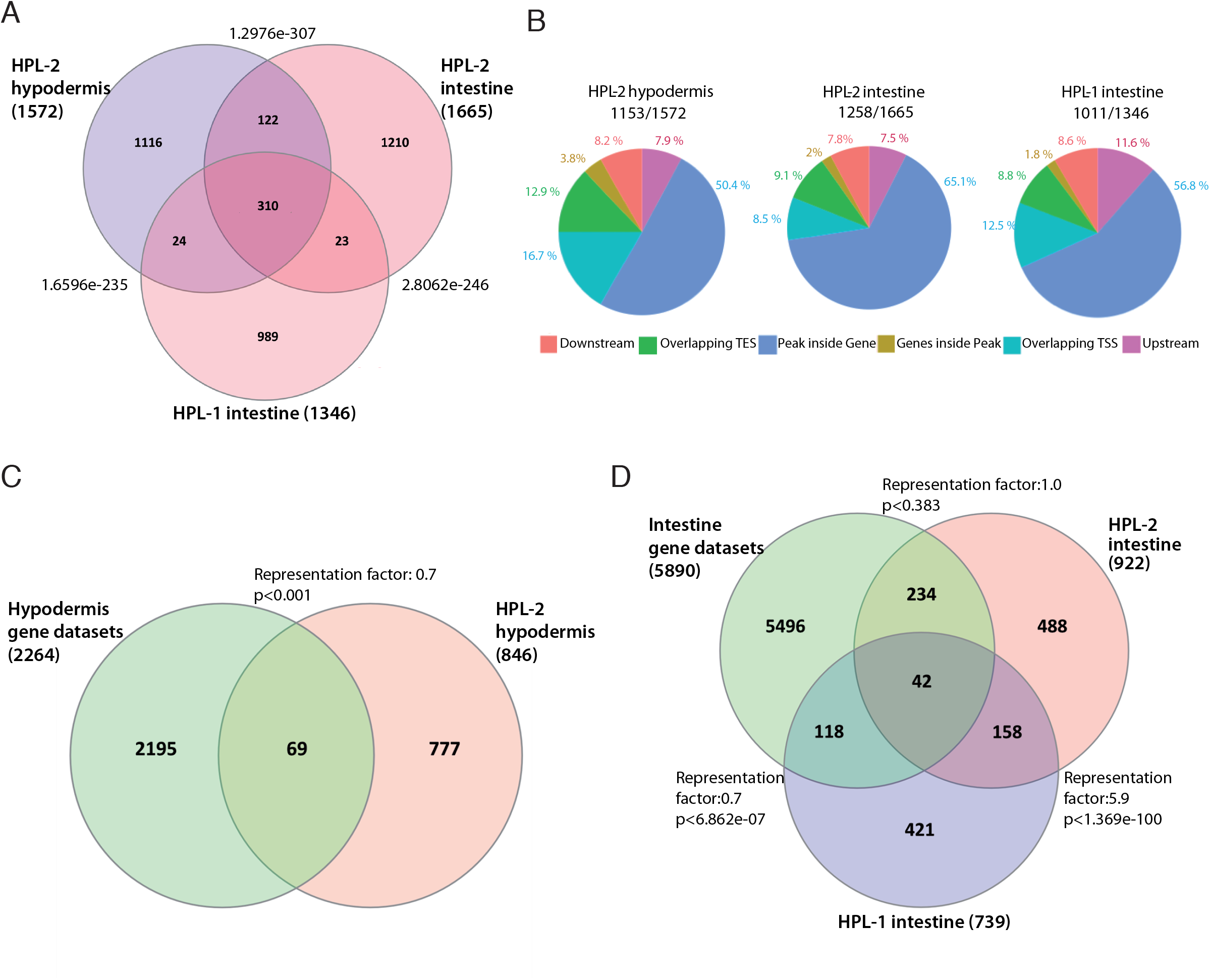
Tissue-specific binding of HP1 proteins and association with gene expression. (A) Venn diagram depicting the overlapping DNA regions bound by HP1 proteins in hypodermis and intestine. The intersection was performed using 10% of fraction reciprocal between genomic regions. Fisher’s exact test p-values are represented. (B) Pie charts showing the percentages of peaks inside various genomic features. Peak assignment was done to the nearest gene whose center coordinate was within 3 kb upstream and 3 kb downstream of the TSS and TES, respectively, of a gene. Ratios above pie charts show the number of peaks assigned to genes relative to the total number of peaks. TSS: Transcription start site; TES: transcription end site. (C) Venn diagram depicting the overlap between HPL-2-bound genes in the hypodermis and genes robustly expressed in this tissue. (D) Venn diagram depicting the overlaps between HPL-1 and HPL-2-bound genes in the intestine and genes expressed in this tissue. Exact hypergeometric probability representation factor and p-values are shown.

Annotation of genomic features for regions bound by HP1 proteins showed that more than 70% of the peaks reside inside protein coding regions (Figure 2B; see peak ratio). Most bound regions are placed within transcribed chromatin, and a notable proportion overlaps with transcription start sites (TSS) or promoter regions (Figure 2B).

In order to know whether genes bound by HP1 proteins are repressed or activated, a comparison with gene datasets from alternative methods of tissue-specific transcriptomic profiling was performed. First, for hypodermal tissue, we selected genes identified in at least three out of four transcriptomic methods (Blazie *et al*. 2017; Kaletsky *et al*. 2018; Katsanos *et al*. 2021; Fragoso-Luna *et al*. 2023). We found a significant underrepresentation of transcribed genes among HPL-2 hypodermal-bound genes (<3% overlap; p < 0.001), suggesting a repressive role for HPL-2 in the hypodermis (Figure 2C; Supplementary File 1; Table S2). We followed a similar procedure with intestinal genes, resulting in an underrepresentation of transcribed genes when compared with HPL-1-bound genes (<3% overlap; p < 6.862e-07) (Figure 2D; Supplementary File 1; Table S2)(Haenni *et al*. 2012; Blazie *et al*. 2017; Kaletsky *et al*. 2018; GÓmez-Saldivar *et al*. 2020; Katsanos *et al*. 2021). In contrast, HPL-2 in the intestine did not show any tendency (Figure 2D). These comparisons suggest a tissue-specific repressive role for HPL-2 and that HPL-1 and HPL-2 act differently in intestinal cells.

Interestingly, when comparing HP1 bound genes across the three datasets, we found that around 60% of genes are unique in each dataset, similar to the proportion of unique binding sites (Figure 3A; compare with Figure 2A; Supplementary File 2). Gene ontology (GO) enrichment analysis for biological processes with genes bound by HPL-2 in the intestine identified only a few and rather broad terms, such as development and anatomical structures (Figure 3B). Because these terms are based on genes primarily expressed in proliferative tissues, their association with HPL-2 in terminally differentiated intestinal cells is concordant with a repressive role of HPL-2. Similarly, gene sets retrieved from either HPL-2 DamID in hypodermal tissue or HPL-1 DamID in the intestine identified only a few enriched GO terms and none of them related to the tissue where the Dam fusion protein was expressed (Figure S6A-B).

**Figure 3.**
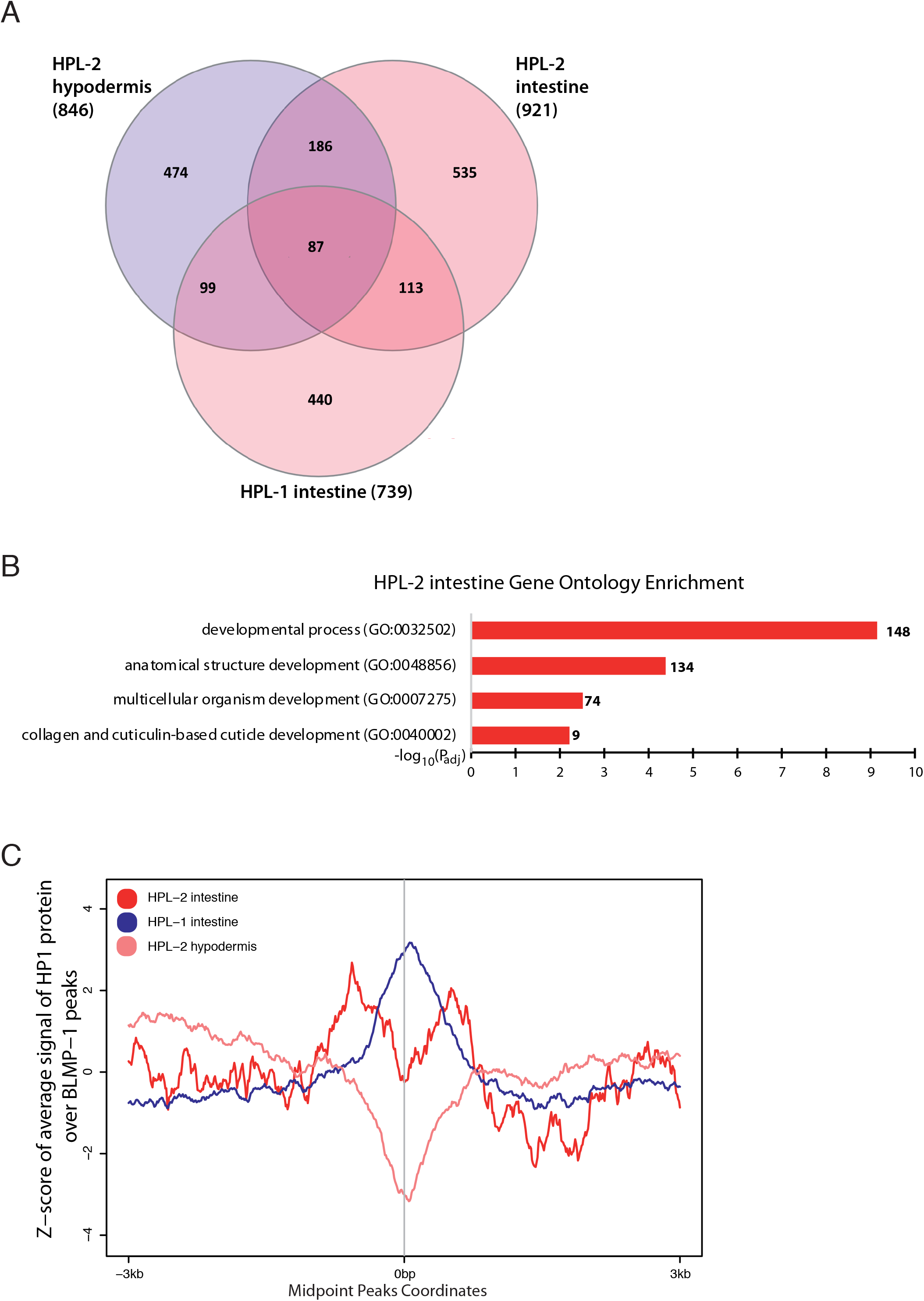
Differential tissue-specific binding of HP1 proteins. (A) Venn diagram depicting the overlapping genes bound by HP1 proteins in the different tissues. Exact hypergeometric probability representation factor and p-values are shown. (B) GO terms associated to Biological process category of genes bound by HPL-2 in the intestine. Number of intersected genes are depicted at the end of the bars. (C) Aggregation plot depicting the Z-score of aggregated average signal of HP1 proteins over 3 kb upstream and 3 kb downstream of BLMP-1 peaks’ midpoint position.

In our attempt to get more insight about new biological functions of HP1 proteins, we compared HP1-bound regions with those bound by different transcription factors at the L1 larval stage (ALR-1, MAB-5, EOR-1, PQM-1, PHA-4, ELT-3, ELT-2, EGL-5, SKN-1, UNC-130, EGL-27 and BLMP-1) (Niu *et al*. 2011). Interestingly, this analysis revealed a significant overlap between the transcription factor BLMP-1/PRDM1 and HPL-1 in the intestine, which is maintained at the L4 stage as well (Figure S6C-D; Table S3)(Stec *et al*. 2021). Around 13% of peaks are common to HPL-1 and BLMP-1, mainly in autosome centers in L1 (Figure S6E). In line with this, the genome-wide average signal of HP1 binding at BLMP-1 peaks showed a strong enrichment for HPL-1 (Figure 3C). Then, we assigned the BLMP1 significant peaks to genes using the same criteria and method employed for HP1 datasets describe here. We found a significant overlap between BLMP-1 and HP1 bound genes, including more than one hundred genes in common (Figure S3F). Collectively, those results strongly suggest that HPL-1 and BLMP-1 could potentially co-regulate specific genes or transcription factors. Knowing that PRDM1, the human BLMP-1 orthologue, was first described as a repressor of beta-interferon (beta-IFN) expression, a tissue-specific co-repression by HPL-1 and BLMP-1 may occur (Keller and Maniatis 1991).

### Only HPL-2 protein is enriched at repetitive elements

It has been previously reported a role for several heterochromatin proteins, among them HPL-2, in maintaining repetitive elements and particular genes silenced, which is crucial for germline and fertility processes (Ashe *et al*. 2012; McMurchy *et al*. 2017). HPL-2 is also enriched at repetitive elements both, in embryos and young adults (Garrigues *et al*. 2015; McMurchy *et al*. 2017). These analyses were performed in whole animals and hence offered no information about tissue-specificity nor the behavior of HPL-1. Therefore, we assessed potential differences in the binding of HPL-1 and HPL-2 to repetitive elements using annotation from the most recent UCSC Genome Browser based on repeat masker that classified 61,527 individual repetitive elements (https://genome.ucsc.edu)(Tarailo-Graovac and Chen 2009). Interestingly, our analysis corroborates the high proportion of overlapping regions between HPL-2 and genomic repetitive elements in both tissues (Figure 4A; Table S4). However, the overlap is reduced and not significant in the case of HPL-1 protein (Figure 4A; Table S4), indicating that HPL-1 is not enriched at repetitive elements.

**Figure 4.**
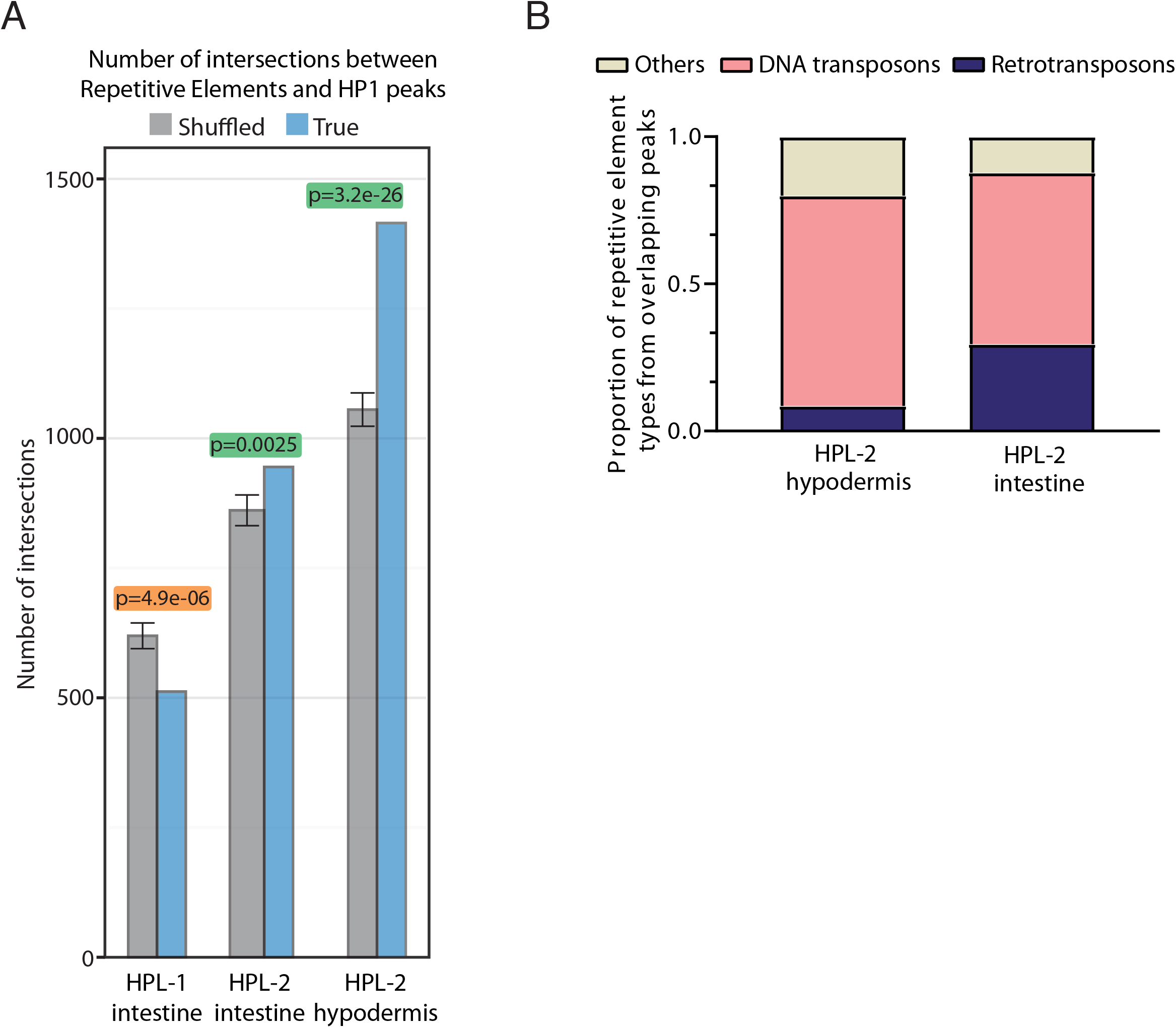
Associationof HP1 proteins to repetitive elements. (A) Number of intersected peaks between HPL-1 and HPL-2 and repetitive elements assessed by Monte Carlo simulation statistical test. Green-shaded p-values: number of overlaps are higher than expected by chance. Orange-shaded p-value: Intersections are lower than expected by chance. (B) Stacked plot depicting the proportion of repetitive element types bound by HPL-2 datasets (See Table S5).

Moreover, the hypodermal tissue exhibited 14% more overlapping regions compared to the intestine for HPL-2 protein, remarking tissue-specific divergence. To further study this difference, we looked at repetitive element types, finding DNA transposon as the most predominant type for both tissues (Figure 4B). Being CELE14B, CELE42 and CELE14A the most frequent DNA transposons families found for both tissues (Table S5). However, more than a quarter of repetitive elements uniquely enriched in the intestinal tissue correspond to retrotransposons, with almost no representation in the hypodermis (Figure 4B). The strongest association reported for HPL-2 in young adults was found with DNA transposons Helitron families, being less abundant compared to other families at the L4 stage analyzed here (Table S5)(McMurchy *et al*. 2017). Nonetheless, for all stages analyzed here and in other studies, DNA transposons are the most representative, suggesting the necessity to maintain these repressed in differentiated tissues.

## Discussion

Here we characterized the genomic binding profile of the two *C. elegans* HP1 protein homologs HPL-1 and HPL-2 in the intestine, as well as the binding profile of HPL-2 in the hypodermis. We showed that HPL-2-bound regions are enriched in distal chromosome arms, as published for embryos and young adults (Garrigues *et al*. 2015; McMurchy *et al*. 2017). In contrast, we showed an unprecedented location of HPL-1 in the center of autosomes.

When assessing the binding to heterochromatic histone marks, we found a strong correlation of HP1 proteins with H3K9me3 and H3K27me3. On the one hand, methylated H3K9 prevents the access of transcription factors to their binding sites in differentiated cells, being crucial for restricting plasticity and maintaining tissue identity. On the other hand, H3K27me3 seems to limit plasticity, acting in a parallel manner (Patel and Hobert 2017). Our data suggest that HP1 proteins act non-redundantly in controlling tissue-specificity in differentiated somatic cells. In agreement with this, cellular plasticity is affected both in single *hpl-1* or *hpl-2* mutants and more strongly upon deletion of both HP1 proteins (Patel and Hobert 2017). Moreover, Evans et al. described some domains decorated by H3K27me marks, considered the most suceptible to be regulated during development (Evans *et al*. 2016). Lastly, in embryos, genes expressed in terminally differentiated tissues were marked specifically by H3K9me3 (Zeller *et al*. 2016). It could be that HP1 proteins contribute to the formation of these domains, as well as controlling tissue-specific genes, affecting chromatin activity and gene expression in a context-dependent manner. In other words, the ability of HP1 proteins to bind concurrently to both repressive histone modifications, reinforce the repression, and could account for developmental gene regulatory networks. Nevertheless, the lack of tissue-specific profiles for histone methylation limits the possibility to interpret in an exhaustive manner the behavior of HPL-1 and −2 in differentiated hypodermal and intestinal tissues in terms of their association to particular histone marks. Recently, single cell analyses have revealed that histone modifications contribute to cell heterogeneity within and between tissues (Carter and Zhao 2021). In addition, H3K9me histone mark deposition is essential to confer tissue-specific gene expression and preserve the integrity of differentiated muscles (Methot *et al*. 2021). Moreover, in mammals, HP1 protein isoforms show complex cell-type and tissue-specific patterns. While HP1 isoforms had the same pattern in the hepatic tissue, patterns where strikingly different in the lymph nodes (Ritou *et al*. 2007). However, our approach is relevant considering the relative contribution of intestinal and hypodermal tissues in worms (Froehlich *et al*. 2021).

HPL-2 showed a strong correlation with H3K9me1 and H3K9me2 in both tissues analyzed, which was not observed with HPL-1. The different preferences of the two *C. elegans* HP1 proteins is reminiscent to the reported association of the three Drosophila homologs, HP1a, HP1b and HP1c to specific classes of chromatin (Schoelz *et al*. 2021) and their different affinities towards H3K9me isoforms (Lee *et al*. 2019). In embryos, HPL-2 closely associated with H3K9me1 and H3K9me2, and less with H3 tri-methylated forms (Garrigues *et al*. 2015). Moreover, more than 20% of regulatory elements were uniquely enriched for H3K9me2 in this stage, mostly tandem or simple repeats type (Zeller *et al*. 2016). This could be in line with the tight association between HPL-2, but not HPL-1, and repetitive elements previously reported in other stages (Garrigues *et al*. 2015; McMurchy *et al*. 2017). Those repetitive elements are mostly represented by DNA transposon families, conversely to retrotransposon families. HPL-2 could bind to H3K9 di-methylated forms to prevent de-repression of repeats and maintain the genome instability during development. The higher proportion of repetitive elements enrichment among HPL-2 peaks in the hypodermis compared to the intestine suggests a specific regulation of genome silencing, highlighting both unique and more general role for HPL-2 during tissue differentiation.

Lastly, the significant underrepresentation of transcribed genes bound by HP1 proteins in either hypodermis or intestine strongly suggests a general repressive role for both HPL-1 and HPL-2. However, the reduced number of peaks (thus genes) shared by all HP1 datasets indicate that they control gene expression in a tissue-specific manner. In fact, in flies, the characterization of HP1a, HP1b, and HP1c protein homologs revealed a tissue-specific pattern and a reduced number of canonical target genes (Schoelz *et al*. 2021). This is likely influenced by histone modifications, but may also be regulated by phosphorylation of HP1 proteins which alter their association with chromatin (Zhao *et al*. 2001; Hiragami-Hamada *et al*. 2011).

Our analysis showed a significant number of genomic regions commonly bound by intestinal HPL-1 and the BLMP-1/PRDM1 transcription factor in two developmental stages. In *C. elegans* BLMP-1 is relevant for the proper development of several tissues, such as, vulva, hypodermis and gonad (Horn *et al*. 2014; Huang *et al*. 2014; Yang *et al*. 2015). PRDM1 was first described as responsible for B cell identity by repressing other transcription factors, thereby controlling indirectly numerous target genes (Shaffer *et al*. 2002). Moreover, during cell plasma differentiation, PRDM1 recruits histone deacetylase HDAC1/2 and lysine demethylase LSD1 chromatin regulators, to repress mature B-cell gene expression pattern (Su *et al*. 2009). In worms, the recruitment of HDACs proteins through the binding of HPL-2 to H3K27me3 represses Hox genes (Jedrusik-Bode 2013). We speculate that

HPL-1 and BLMP-1 could act in parallel to maintain silent genomic regions crucial for cell identity. As occurs in plasma cells, BLMP-1 may co-regulate with HPL-1 and other epigenetic factors the repression of specific genomic regions in order to control cell fate. The specific mechanism behind remains elusive, but a recent study proposed that BLMP-1 regulates chromatin accessibility during development (Stec *et al*. 2021).

In summary, this tissue-specific study of HP1 proteins revealed important differences that highlight their relevance as heterochromatic markers for cell identity during development. This potential restriction of plasticity accompanied by changes in other crucial heterochromatin marks, such as histones modifications, open new questions about the role of chromatin modifiers in this complex process. Lastly, we revealed unique features for the underexplored HPL-1 homolog in *C. elegans*, which encourage further studies to uncover its biological roles.

## MATERIAL AND METHODS

### *C. elegans* strains and maintenance

Maintenance of *C. elegans* strains were done conforming to standard protocols on Nematode Growth Medium (NGM) plates, grown in a monoxenic lawn of *Escherichia coli* OP50 at 20°C (Stiernagle 2006). For DamID experiments, a lawn of *E. coli* GM119 dam(-) strain was used to grow the worms (Arraj and Marinus 1983). The complete list of strains used in this study is available in Table S6.

### Molecular cloning

To generate the Dam::HPL-1 expression construct pBN494 (P*hsp-16*.*41::FRT::mCherry::his-58::FRT::Dam::hpl-1*) the *hpl-1* CDS was PCR amplified from genomic DNA and inserted in the pCRII cloning vector (Invitrogen). Next, MYC-flanked Dam was inserted into the *Eco*RI site in the first exon of *hpl-1*. Finally, Dam::*hpl-1* was excised with *Xho*I + *Spe*I and inserted into pBN488 (Fragoso-Luna *et al*. 2023) digested with *Xho*I + *Nhe*I. To generate the Dam::HPL-2 expression construct pBN495 (P*hsp-16*.*41::FRT::mCherry::his-58::FRT::Dam::hpl-2a*) the *hpl-2a* CDS was PCR amplified from genomic DNA and inserted in the pCRII cloning vector. Next, MYC-flanked Dam was inserted into the *Bam*HI site in the third exon of *hpl-2a*. Finally, Dam::*hpl-2a* was excised with *Sal*I + *Spe*I and inserted into pBN488 digested with *Xho*I + *Nhe*I. This insertion results in the specific tagging of the *hpl-2a* isoform. Designs were based on previous constructs for multicopy (Couteau *et al*. 2002) and endogenous (Patel and Hobert 2017) tagging of HPL-1/2.

### Mos Single Copy Insertion (MosSCI)

Single copy integration transgenic strains were made by microinjection into strain EG4322 (ChrII) with *unc−119(+)* recombination plasmids together with individual transgenes (pBN494 and pBN495) at 25 ng/μl, Mos1 transposase (P*eft-3*::transposase) at 50 ng/μl and three red co-injection markers pCFJ90 (P*myo-2::mcherry*) at 2,5 ng/μl, pCFJ104 (P*myo−3::mcherry*) at 5 ng/μl and pBN1 (P*lmn-1::mcherry::his−58*) at 10 ng/μl (FrØkjÆr-Jensen *et al*. 2012; Dobrzynska *et al*. 2016). Plasmids were microinjected into one of the gonad arms of egg-laying and locomotion defective *unc-119* young adult worms. Injected worms were individually transferred to NGM petri dishes and placed at 25°C. Successful integrants were spotted after 7−14 days based on wild type mobility and absence of co−injection markers. Description of plasmids can be found in Table S7.

### DamID-seq protocol

DamID experiments were performed as described in (de la Cruz Ruiz *et al*. 2022). Strains were synchronized by standard hypochlorite treatment and grown until the L4 stage. Optimal number of 14x cycles were selected for PCR amplification of methylated GATC sites.

### DamlD-seq data processing and visualization

The quality control of the DamID technique and the calculation of chromosome association profiles at 100 kb level for all HP1 samples were performed by damid.seq.r pipeline version 0.1.3 in Rstudio (R version 3.4.3) (available at github.com/damidseq/rDamlDSeq) (Sharma *et al*. 2016). From FastQ files, the pipeline detects and maps reads containing the DamID adapters (adapt.seq=“CGCGGCCGAG”) and match them to genomic regions flanked by GATCs sites (restr.seq = “GATC”) to obtain Bam Files. After mapping with the *C. elegans* genome release “BSgenome.celegans.ucSc.ce11”, the log2 ratio HPL::Dam/GFP::Dam across two replicas was integrated to plot the relative read counts at 100 kb bin along the genome for all chromosome autosomes. Pairwise Wilcoxon rank test values were calculated in R. Coordinates of arms as defined in (GonzÁlez-Aguilera *et al*. 2014; Sharma *et al*. 2016). Read numbers for all datasets are summarized in Table S8.

For the visualization using UCSC Genome Browser, the Bam files obtained in the previous pipeline (damid.seq.r) were used to generate the normalized log2 HPL::Dam/GFP::Dam ratio scores per GATC genome fragments, using perl script damidseq_pipeline v1.4.5 (Marshall and Brand 2015) (available at https://github.com/owenjm/damidseq_pipeline). The pipeline integrates a Bowtie 2 v2.3.4 (Langmead and Salzberg 2012) calling to map reads on *C. elegans* bowtie indexes from assembly WBcel235 (available in Illumina iGenomes webpage; identical to ce11). In addition, it includes also Samtools v1.9 (Li *et al*. 2009) for alignment guidance, and a GFF file containing all GATC sites in the *C. elegans* genome. Last GATC coordinates file was built by gatc.track.maker.pl tool provided by the same pipeline script using WBcel235 FASTA file for *C. elegans* (available at https://ensembl.org/Caenorhabditis_elegans/Info/Index. Final bedgraph files for each replica was averaged using average_tracks perl script included in damid_misc (available at https://github.com/owenjm/damid_misc). For comparison of chromosome IV arms versus centers, we used the following border coordinates: chrIV 4,790,000 & 12,610,000.

For the heatmap depicting Pearson correlations between Histone marks datasets we did a preprocessing to average the replicas in bedgraph/wig format using WiggleTools (Zerbino *et al*. 2014). Then, a fixed bin size was selected to standardized all datasets by “bedgraph_to_wig.pl” tool (created by Sebastien Vigneau in Alexander Gimelbrant lab; available at https://gist.github.com/svigneau/8846527). After converting wig files to bigwig format, we lifted-over the bigwig to equivalent genome version using Crossmap tool (Zhao *et al*. 2014). Finally, the matrix and heatmap were generated by “multiBigwigSummary” and “plotCorrelation” tools inside Deeptools (RamÍrez *et al*. 2016). Sources for datasets listed in Table S9.

### Peak calling and analysis

The selection of statistically significant peaks for every HP1 sample was done by find_peaks perl script (available at https://github.com/owenjm/find_peaks) with a false discovery rate (FDR) lower than 0.05 using as input averaged bedgraph files containing normalized log2 HPL::Dam/GFP::Dam ratio scores per GATC fragment.

Intersection between peaks from different datasets and statistical calculation of Fisher exact test, were performed by “Intersect” and “Fisher” tools included in Bedtools suit (Quinlan and Hall 2010). We include r and f=0.1 options, default mode for the rest. Default options were used for the characterization of repetitive element types. When needed, a lift-over between genome ce10, ce6 and ce11 was performed by “liftOver” tool (available in https://genome.ucsc.edu/cgi-bin/hgLiftOver). Pairwise association for peaks comparison was performed by intervene tool, using aforementioned bedtools options (Khan and Mathelier 2017). Heatmaps and Stacked plots for peaks were generated by GraphPad Prism 8 (www.graphpad.com). The coordinates for comparison between chromosome autosome arms and center were used as described in (Garrigues *et al*. 2015).

A statistical method based on Monte Carlo simulation was applied to plot intersected peaks in Figure S5B and C and Figure S6E and Figure 4A (FerrÉ *et al*. 2019).

### Gene assignment and analysis

The statistically significant peaks selected were assigned to genes using UROPA (Kondili *et al*. 2017) as a web tool (available at http://loosolab.mpi-bn.mpg.de/UROPA_GUI/) with Caenorhabditis_elegans.WBcel235.107.gtf (from http://ensembl.org/Caenorhabditis_elegans/Info/Index) for genome protein coding annotation. Peaks were determined to genes on any strand when their center coordinate was localized up to 3 kb upstream of a gene start site or 3 kb downstream of the gene end site. We used UROPA tool to annotate genes from significant peaks of BLMP-1 dataset. Venn diagram overlapping gene datasets was performed using “Interactivenn” webtool (available in http://www.interactivenn.net/index.html) (Heberle *et al*. 2015). The statistical significance by hypergeometric distribution was done with http://nemates.org/MA/progs/overlap_stats.htmlwebtool.

The Gene Ontology analysis was done using unique genes from HP1 datasets, by g:Profiler webtool using default options (Raudvere *et al*. 2019).

### Aggregation plot of signal localization

All aggregation plots were generated using Seqplots, as a GUI application (Stempor and Ahringer 2016). Plots represent the average signal for every 10 bp bins of depicted histone marks or HP1 datasets over the midpoint position of significant peaks depicted.

### Data Availability

Plasmids and strains can be requested from the authors. Supplemental files are available at FigShare. Supplementary File 1 contains the overlap between HP1 bound genes with tissue-specific expressed genes. Supplementary File 2 contains the overlap between tissue-specific genes bound by HP1 proteins. Datasets generated in this paper are available at deposited at Gene Expression Omnibus (GEO) accession GSE222056 (https://www.ncbi.nlm.nih.gov/geo/).

## Acknowledgements

Very special thanks to Ildefonso Cases from the CABD Bioinformatics Facility for support with data analysis.

## Author contributions

P.dl.C.R., P.A and M.A.-S. designed the experiments; P.dl.C.R. and M.J.R.-P. carried out experiments, P.dl.C.R. analyzed the data; P.dl.C.R., P.A and M.A.-S. interpreted results and wrote the manuscript. All authors read, commented and approved the final manuscript.

---

**Figure S1.**
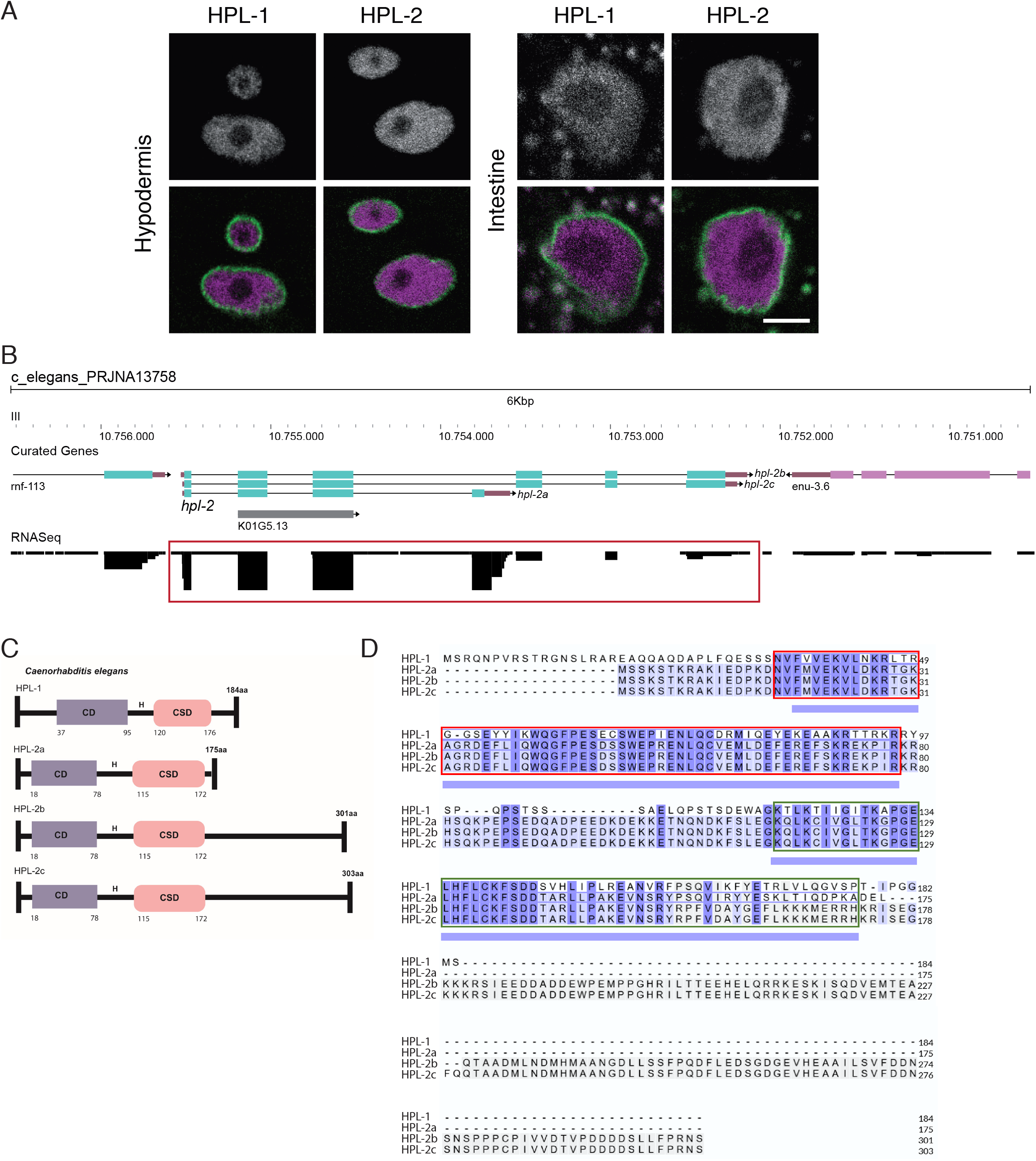
Expression, structure and sequence of *C. elegans* HP1 proteins. (A) Representative microscopy images of mKate2::HPL-1 and mKate2::HPL-2 nuclear expression in the hypodermis (left panel) and intestine (right panel). mKate is shown in magenta, nuclear membrane is labelled by GFP::MEL-28. Bar: 5μm. (B) RNAseq expression data of the different *hpl-2* isoforms (framed in red) from WormBase Genome Browser. Neighboring genes are also shown. (C) Schematic representation of Chromodomain (CD), Chromo Shadow Domain (CSD) and Hinge region (H) in HPL-1 unique isoform and HPL-2 a, b and c isoforms. (D) Protein sequence alignment of HPL-1 (Uniprot ID: G5EET5), HPL-2a (Uniprot ID: GSEDE2-1), HPL-2b (Uniprot ID: G5EDE2-2) and HPL-2c (Uniprot ID: G5EDE2-3). CD depicted by red chart. CSD indicated in green chart. Identical amino acids are highlighted in dark blue. Alignment performed by https://www.uniprot.org.

**Figure S2.**
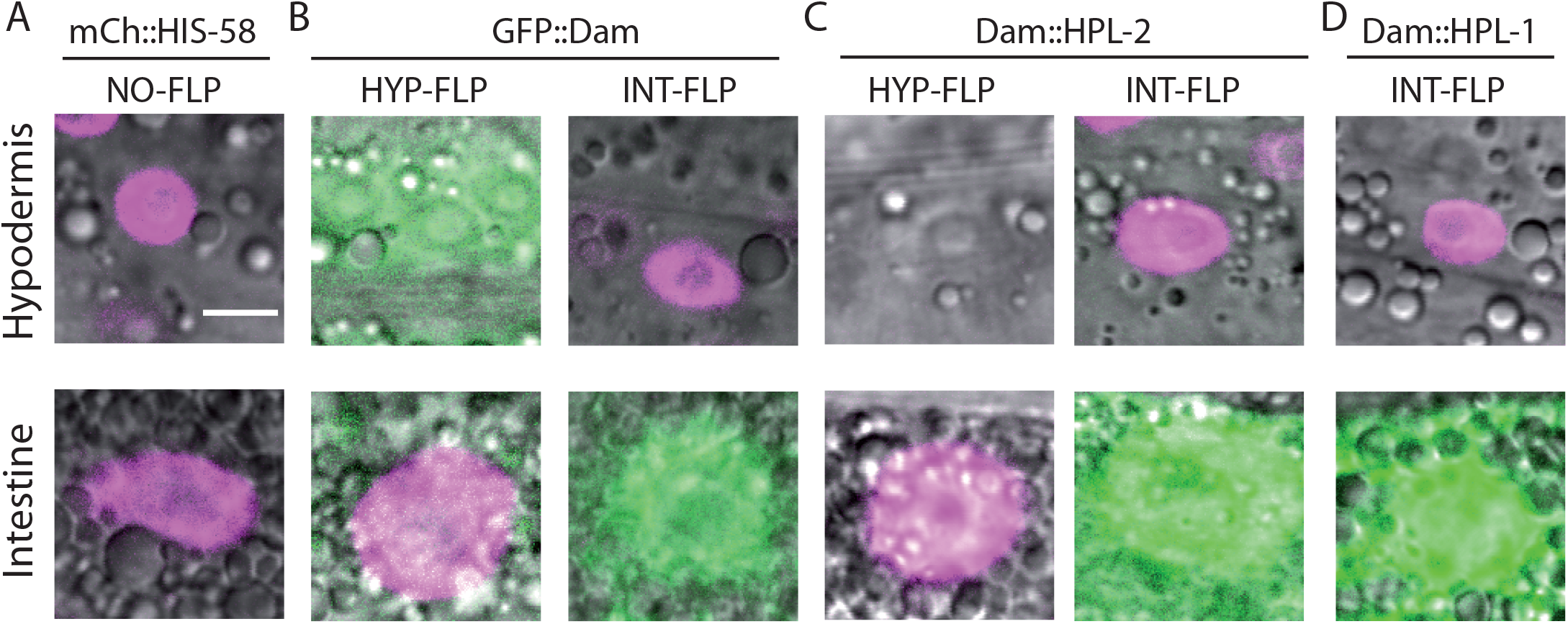
Tissue specificity of the DamID system in hypodermis and intestine. (A) In the absence of Flipase (NO-FLP), the mCherry::*his-58* (mCh::HIS-58) cassette is expressed in all tissues (shown in magenta). (B) GFP::Dam control strains. FLP expression in the hypodermis (HYP-FLP, left panels) or the intestine (INT-FLP, right panels) results in the excision of the mCherry::*his-58* cassette in that tissue and expression of the GFP::Dam fusion protein (shown in green). Expression of mCh::HIS-58 can be observed in the nuclei where FLP is absent. (C-D) Strains used for tissue-specific HPL-1/2 DamID. (C) Expression of FLP in hypodermis (HYP-FLP, left panels) or intestine (INT-FLP, right panels) results in the expression of Dam::HPL-2 in the tissue of interest after excision of the mCherry::*his-58*8 cassette. mCh::HIS-58 can be seen in the other tissue. (D) Dam::HPL-1 expression in the intestine is driven by intestinal FLP (INT-FLP) expression, resulting in the absence of mCh::HIS-58 specifically in that tissue. Note that in B-D INT-FLP is co-expressed with soluble mNeonGreen, which results in green intestinal nuclei and cytoplasm. Bar: 5 μm.

**Figure S3.**
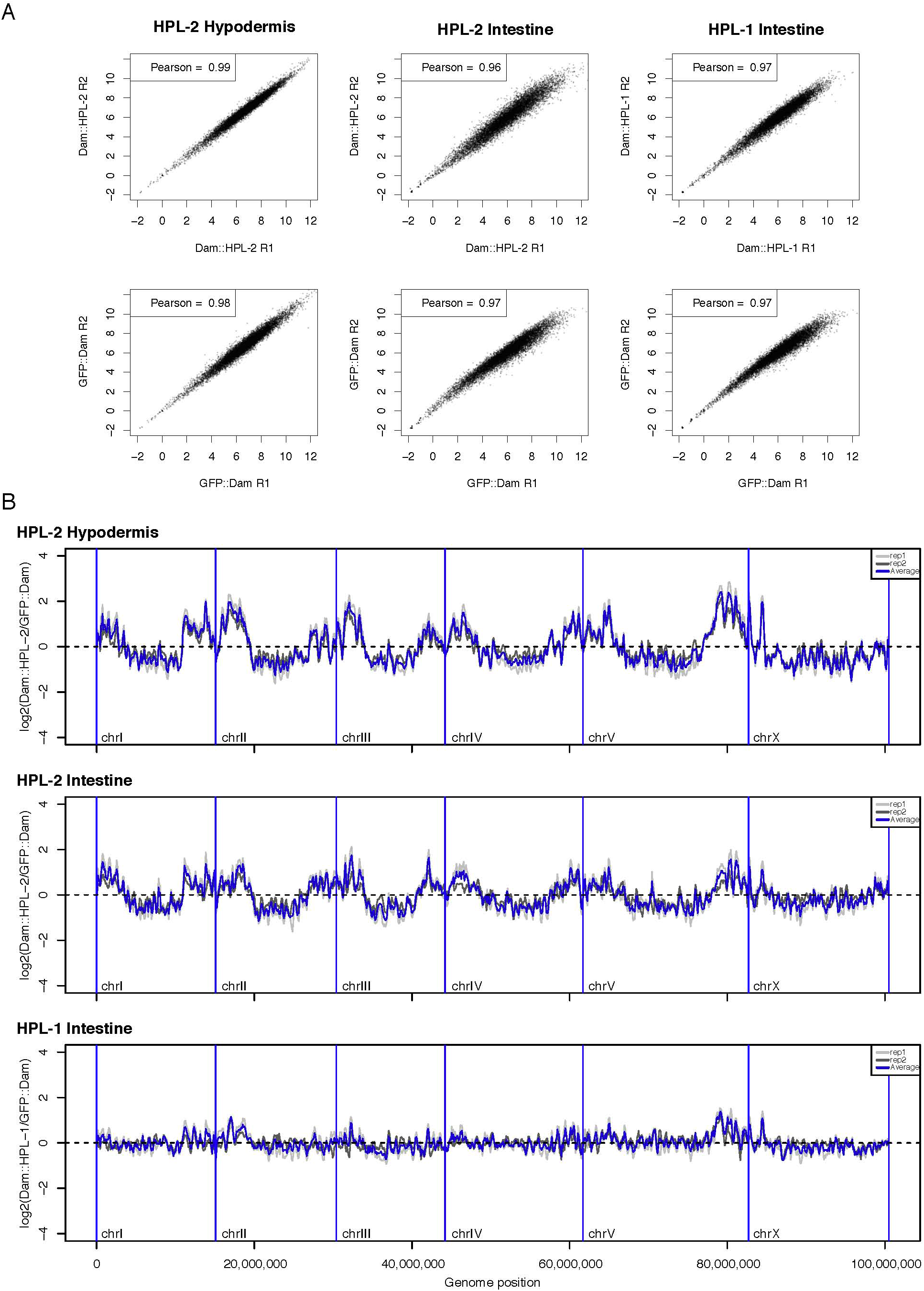
High correlation between DamID replicas. (A) Absolute read numbers per 10kb bin were transformed using the regularized log (rlog) function of the DESeq2 R package (version 1.36.0; (LOVE et al. 2014)). Pearson correlation values for each pair of transformed replicas are indicated in upper left corners. (B) Genome-wide log2(Dam::HPL/GFP: :Dam) profiles at I 00kb bin level. Individual replicas are represented in different tones of grey; average values are represented in blue. To facilitate visualization of the entire genome, the lines represent the rolling mean across 3 bins.

**Figure S4.**
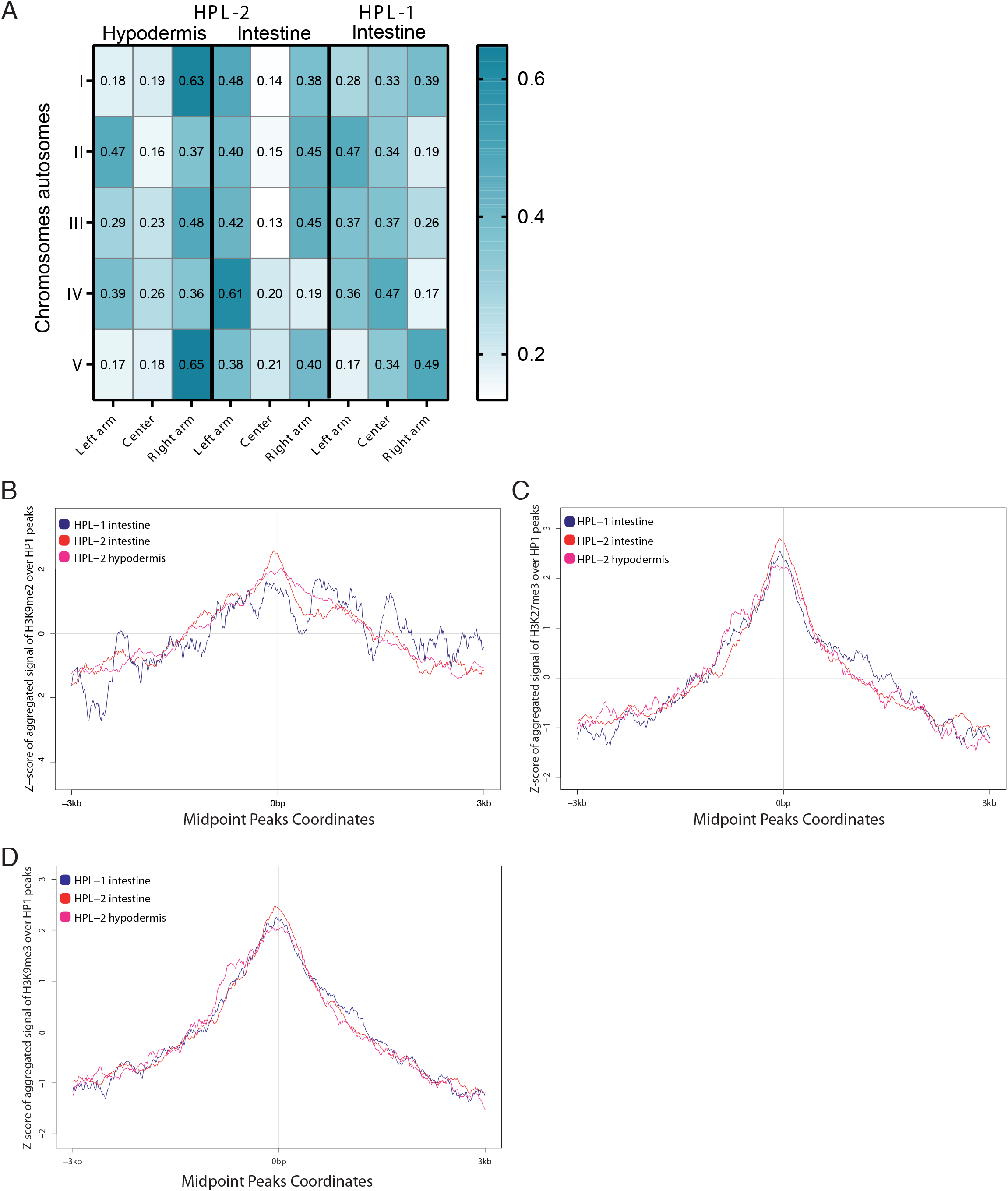
Tissue-specific chromatin binding of HPL-1 and HPL-2 and their association to histone marks. (A) Heatmap depicting the proportion of HP1 peaks distributed by chromosomes autosomes distinguishing between arms and center. Chromosomes autosomes coordinates based on (GARRIGUES et al. 2015). (B-D) Aggregation plots depicting the average signal tracks for H3K9me2 (B), H3K27me3 (C) and H3K9me3 (D) histone datasets over HP1 peaks. Center position represents the midpoint location of peaks. Regions up to 3 kb upstream and 3 kb downstream of the midpoint in 10 bp bins are shown.

**Figure S5.**
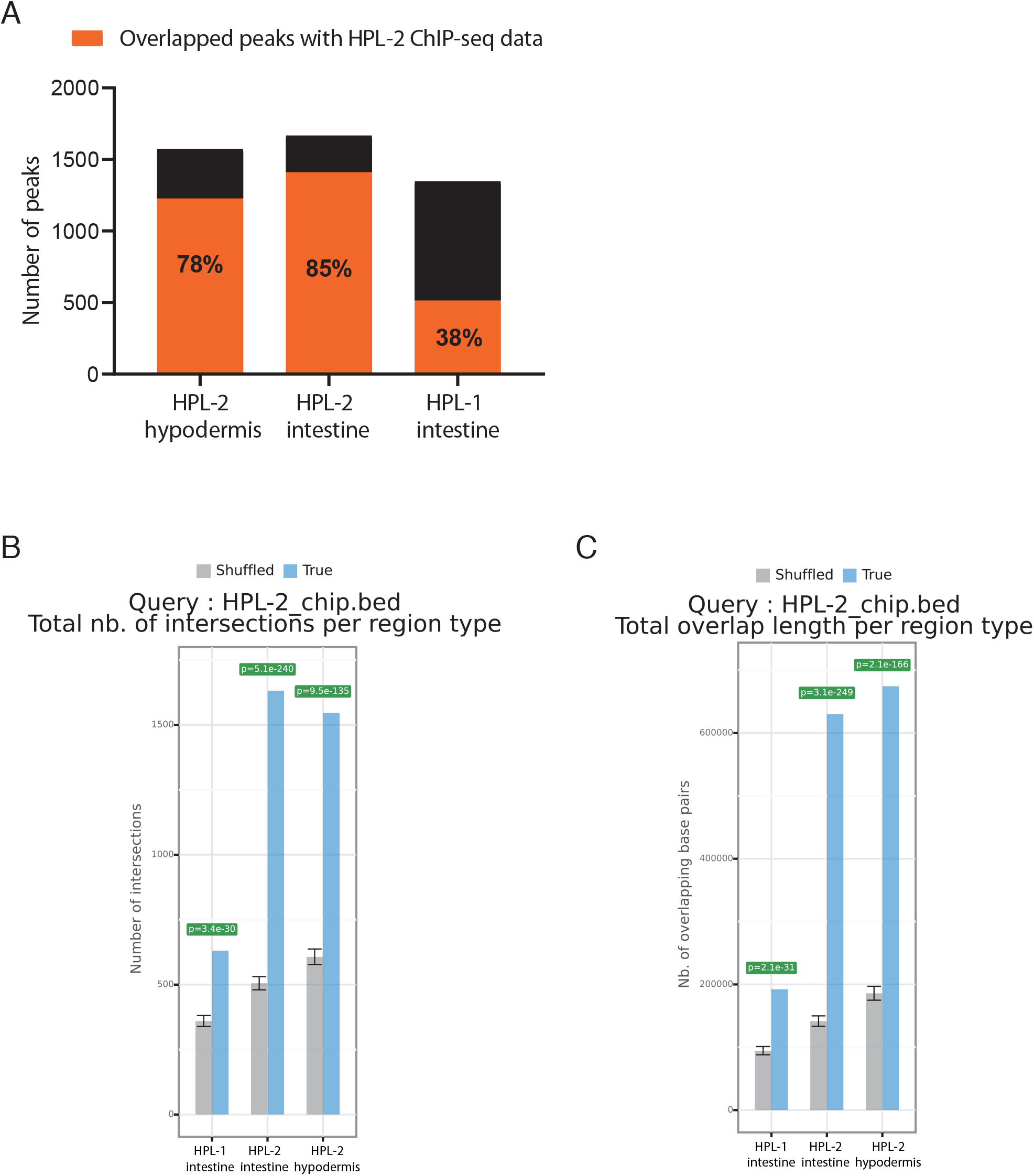
Overlapping of HP1 DamID and HPL-2 ChIP datasets. (A) Stacked plot depicting the number of overlapped peaks between HP1 datasets indicated and a published HPL-2 ChIP dataset. Numbers inside orange bars represent the proportion in percentage of overlapped peaks respect of total HP1 peaks number. (B-C) Statistics for overlapped peaks between HP1and HPL-2 ChIP-seq using a Monte Carlo simulation, based on mathematical distributions of region (B) and inter-region lengths (C), according to negative binomial model of the total overlap length. Hpl-2_hyp: HPL-2 hypodermis; hpl-2_int: HPL-2 intestine; hpl-1_int: HPL-1 intestine.

**Figure S6.**
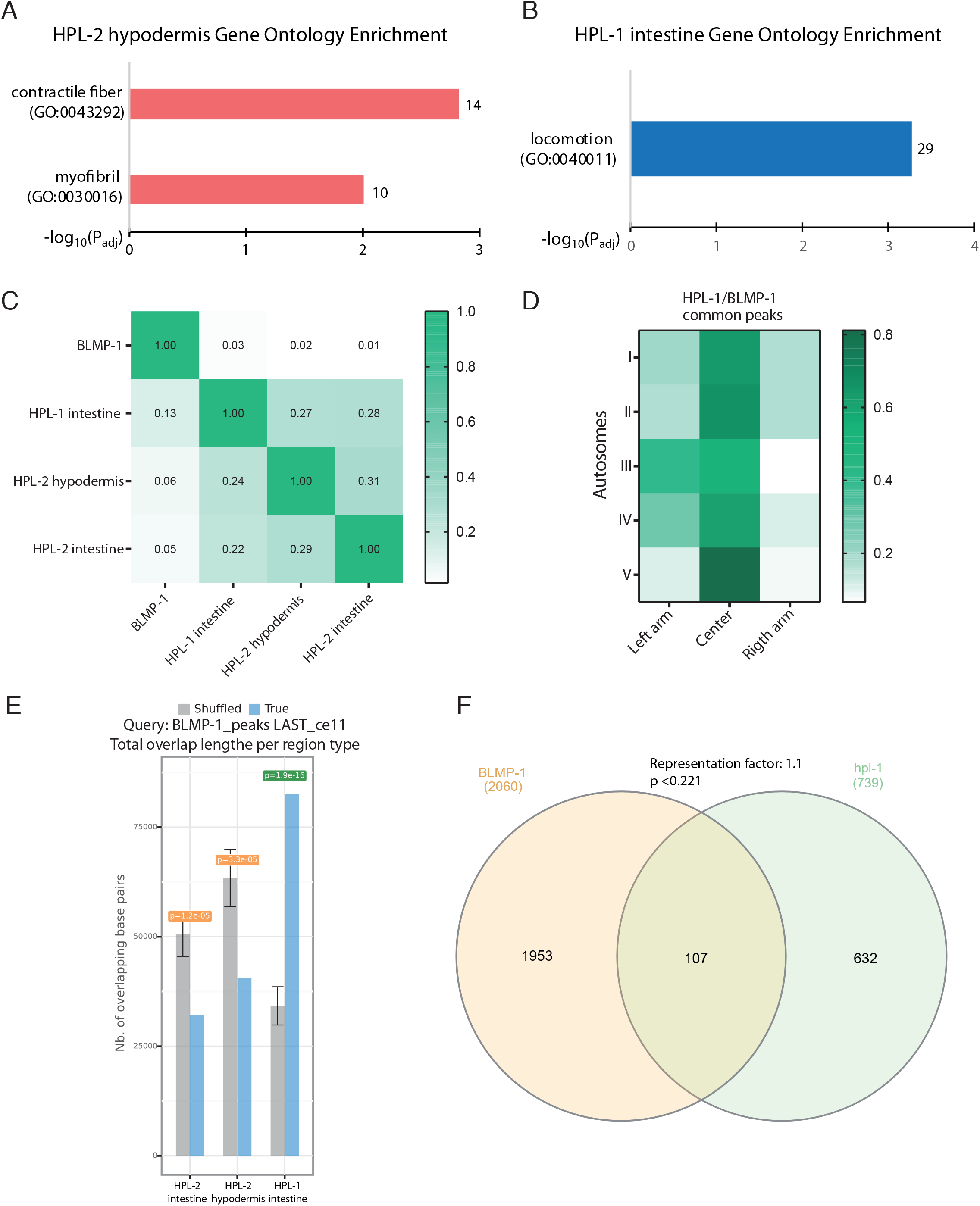
Differential tissue-specific binding of HP1 proteins. (A-B) GO terms associated to cellular component category of unique genes bound by HPL-2 in the hypodermal tissue (A) and by HPL-1 in the intestine (B). Number of intersected genes depicted in right side bars. (C) Heatmap depicting proportion of overlapped peaks between all datasets indicated. The number of BLMP-1 peaks is much higher than the number of HPL-1/2 peaks. Hence, the correlation values when querying with BLMP-1 peaks against HPL-1/2 peaks (top row) are lower than when querying with HPL-1/2 peaks against BLMP-1 peaks (left column). (D) Heatmap describing chromosome autosome location of overlapped peaks between BLMP-1 and HPL-1. (E) Number of intersected peaks between HP1 datasets and BLMP-1 assessed by Monte Carlo simulation statistical test. (F) Venn diagram indicating the number of overlapping genes between HPL-1 in the intestine and BLMP-1. Exact hypergeometric probability representation factor and p-values are shown.

## Supplementary Tables

**Table S1.**
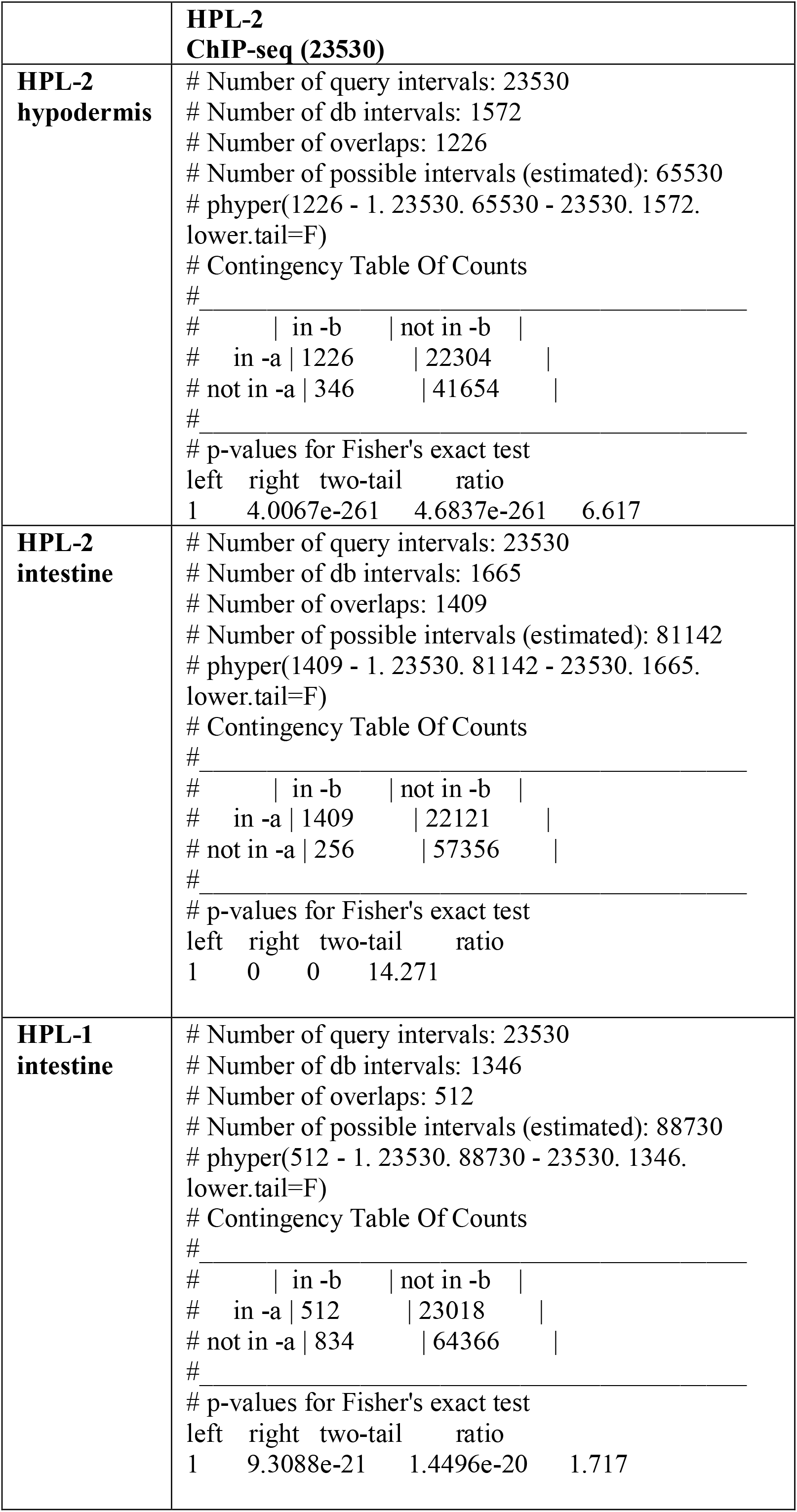
Number of intersections with HPL-2 ChIP-seq genomic regions and Fisher’s exact test p-values.

**Table S2.** Hypergeometric statistical test between gene sets depicted in the table.

|  | Hypodermal well-expressed genes (2264) | Intestinal well-expressed genes (5890) | HPL-2 intestine (922) | HPL-1 intestine (739) |
| --- | --- | --- | --- | --- |
| HPL-2 hypodermis (845) | Representation factor: 0.7<br>$p < 0.001$ | - | | |
| HPL-2 intestine (922) | - | Representation factor: 1.0<br>$p < 0.383$ | - | Representation factor: 5.9<br>$p < 1.369e-100$ |
| HPL-1 intestine (739) | - | Representation factor: 0.7<br>$p < 6.862e-07$ | Representation factor: 5.9<br>$p < 1.369e-100$ | - |

**Table S3.** Number of overlapped peaks between HP1 and BLMP-1 ChIP-seq data and Fisher’s exact test p-value.

| Samples (significant peaks) | Number of intersections with **BLMP-1 peaks (5898)** | Fisher's exact test p-values |
| --- | --- | --- |
| **HPL-1 intestine (1346)** | 169 | two-tail ratio<br>1.8565e-09 1.701 |
| Samples (significant peaks) | Number of intersections with **L4 BLMP-1 peaks (549)** | Fisher's exact test p-values |
| **HPL-1 intestine (1346)** | 26 | two-tail ratio<br>0.013515 1.701 |
| **HPL-2 intestine (1665)** | 34 | two-tail ratio<br>0.0041704 1.726 |
| **HPL-2 hypodermis (1572)** | 14 | two-tail ratio<br>0.097535 0.630 |

**Table S4.** Number of overlapped regions and p-values from Monte Carlo Simulation between HP1 significant peaks and repetitive element coordinates annotation from UCSC based on repeat masker (ce11).

| Samples (significant peaks) | Number of true intersections | p-values |
| --- | --- | --- |
| **HPL-2 hypodermis (1572)** | 1415 | 3.192e-26 |
| **HPL-2 intestine (1665)** | 945 | 0.002534 |
| **HPL-1 intestine (1346)** | 512 | 4.887e-06 |

**Table S5.** Top 20 overlapped regions between HP1 and repetitive elements.

|  |  |  |
| --- | --- | --- |
| **HPL-2 hypodermis** |  |  |
| Family | Type | Number of repeats |
| CELE14B | DNA | 282 |
| CELE42 | DNA | 152 |
| CELE14A | DNA | 92 |
| CELE1 | DNA | 75 |
| PALTA5_CE | DNA | 61 |
| GA-rich | Low_complexity | 48 |
| A-rich | Low_complexity | 46 |
| CeRep5 | Unknown | 44 |
| TIR9TA1B_CE | DNA | 40 |
| CELE2 | DNA | 38 |
| TIR9TA1_CE | DNA | 36 |
| Helitron2_CE | RC | 32 |
| MINISAT1_CE | Satellite | 30 |
| HelitronY1A_CE | RC | 27 |
| HAT1_CE | DNA | 25 |
| CELE45 | SINE | 24 |
| CELE4 | DNA | 23 |
| RC123 | Satellite | 21 |
| PALTTAA2_CE | DNA | 20 |
| CeRep3 | Unknown | 18 |
| **HPL-2 intestine** |  |  |
| CELE14B | DNA | 193 |
| CELE42 | DNA | 112 |
| CELE14A | DNA | 58 |
| TIR9TA1B_CE | DNA | 48 |
| CELE45 | SINE | 42 |
| TIR9TA1_CE | DNA | 40 |
| PALTA5_CE | DNA | 32 |
| GA-rich | Low_complexity | 28 |
| A-rich | Low_complexity | 26 |
| MINISAT1_CE | Satellite | 26 |
| PALTTAA3_CE | DNA | 12 |
| CeRep5 | Unknown | 12 |
| CER12-I_CE | LTR | 12 |
| PALTTAA1_CE | DNA | 11 |
| Vingi-2_CE | LINE | 11 |
| LR9A | Unknown | 11 |
| HelitronY1_CE | RC | 10 |
| CER11-I_CE | LTR | 9 |
| TIR9TA1C_CE | DNA | 8 |
| CELE4 | DNA | 8 |

**Table S6.**
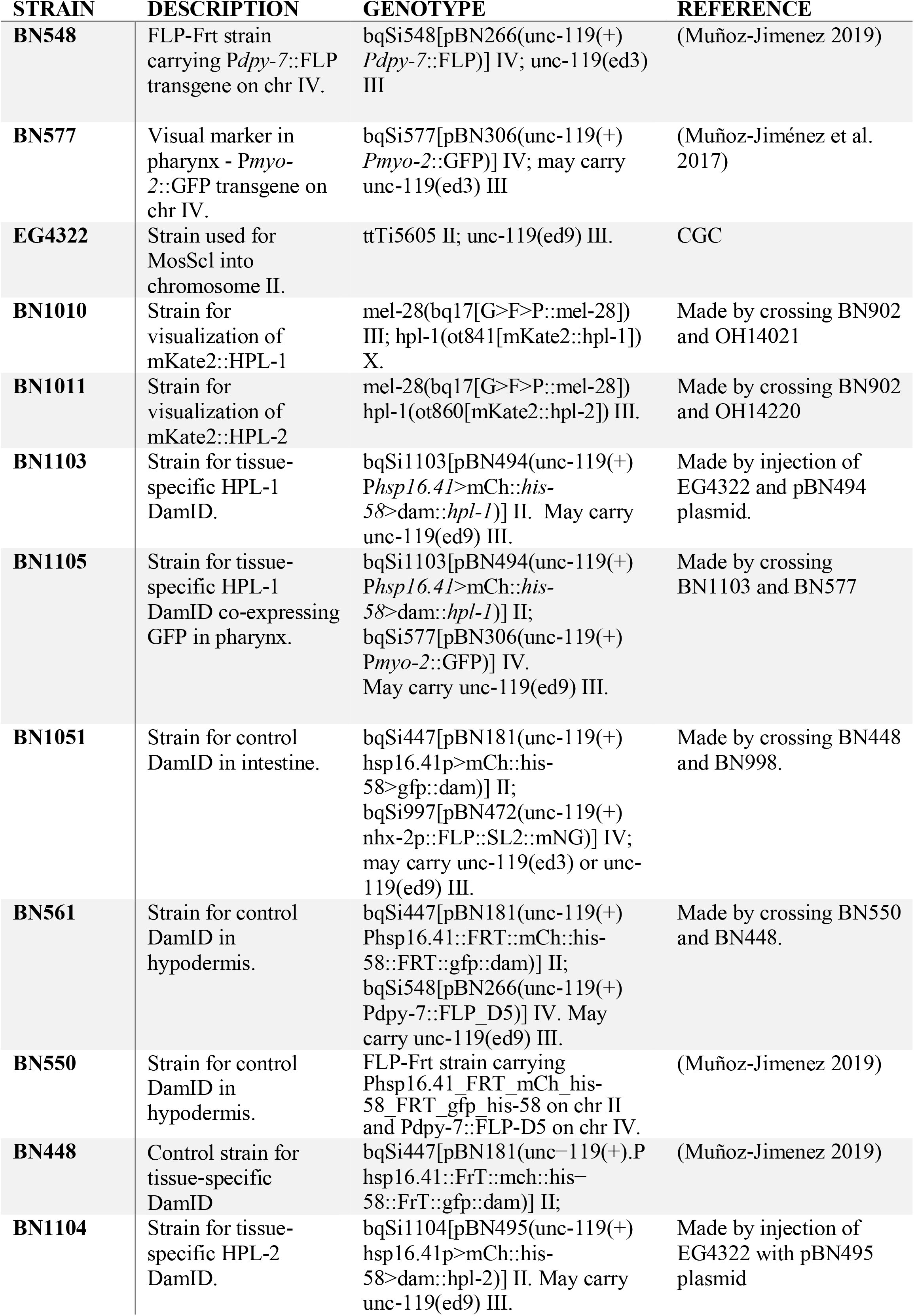
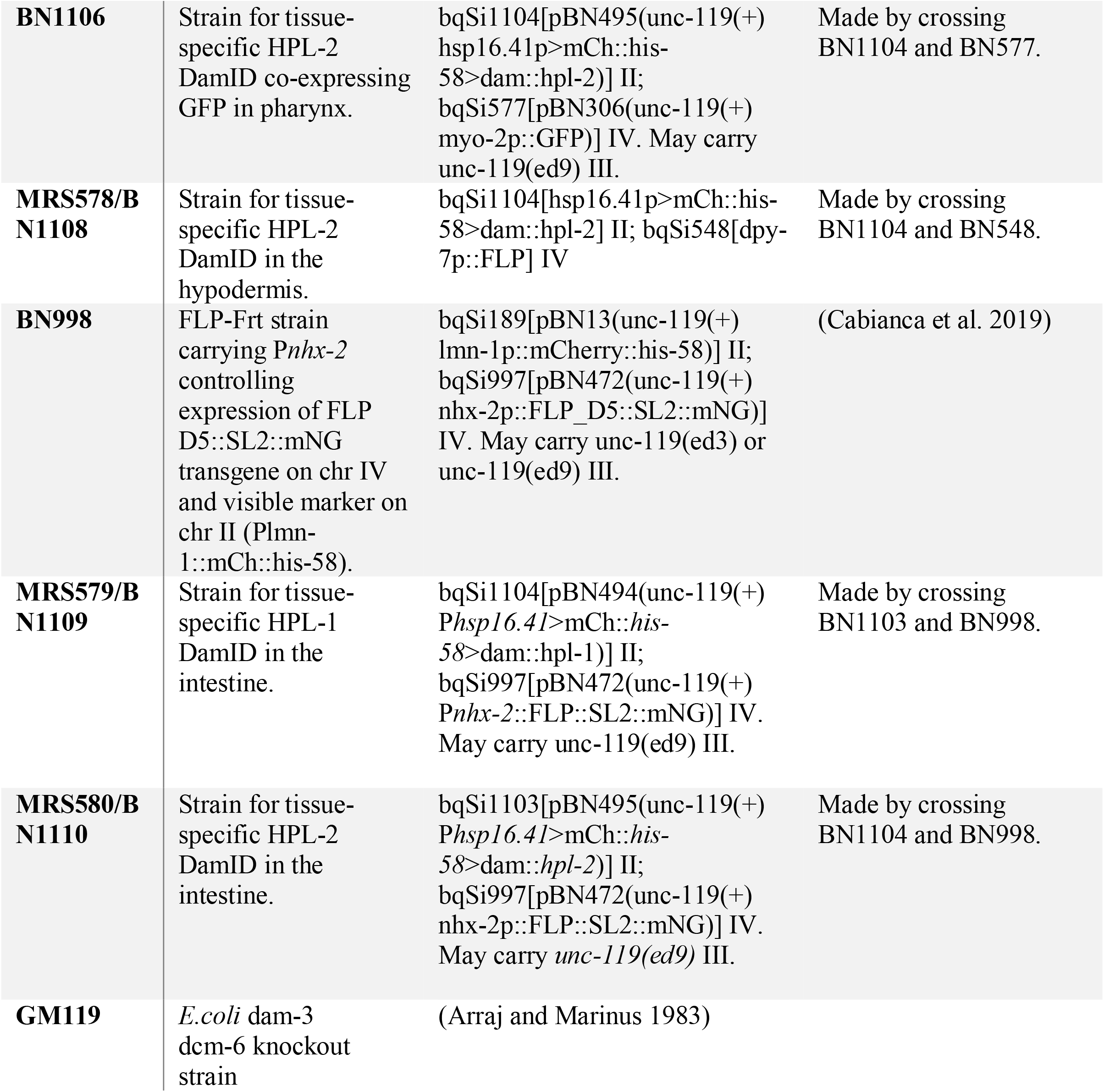
List of *C*.*elegans* and bacterial strains used.

**Table S7.** List of plasmids used.

| Plasmid name | Description | Reference |
| --- | --- | --- |
| **pBN494** | *Phsp-16.41*::FRT::mCh::his-58::FRT::hpl-1::Dam Chr II | This publication |
| **pBN495** | *Phsp-16.41*::FRT::mCh::his-58::FRT::hpl-2::Dam Chr II | This publication |

**Table S8.** Quality controls metrics of sequencing libraries from DamID-seq data. Abbreviations hyp: hypodermis; int:intestine

| Sample | Raw reads | Cut reads | Mapped reads | GATC reads (% from Raw reads) |
| --- | --- | --- | --- | --- |
| GFP::Dam hyp (BN561) I | 15,773,728 | 6,542,411 | 2,872,416 | 2,862,349 (18,15%) |
| GFP::Dam hyp (BN561) II | 9,247,899 | 3,599,466 | 1,304,548 | 1,286,460 (13,91%) |
| GFP::Dam int (BN1051) I | 18,066,270 | 7,418,519 | 3,660,999 | 3,655,161 (20,23%) |
| GFP::Dam int (BN1051) II | 4,573,067 | 2,080,559 | 1,338,144 | 1,336,648 (29,23%) |
| Dam::hpl-2 hyp (MRS578) I | 14,194,267 | 6,551,959 | 4,162,172 | 4,154,179 (29,27%) |
| Dam::hpl-2 hyp (MRS578) II | 8,507,806 | 3,473,687 | 1,594,499 | 1,562,681 (18,37%) |
| Dam::hpl-1 int (MRS579) I | 20,901,460 | 9,376,852 | 1,560,420 | 1,541,018 (39,23%) |
| Dam::hpl-1 int (MRS579) II | 8,061,225 | 3,861,200 | 696,028 | 694,399 (8,61%) |
| Dam::hpl-2 int (MRS580) I | 18,337,259 | 8,265,167 | 4,253,333 | 4,232,349 (23,08%) |
| Dam::hpl-2 int (MRS580) II | 1,583,607 | 747,426 | 527,174 | 526,083 (33,22%) |

**Table S9.** Description and references of Dataset used for comparison.

| Sample name | Dataset (GEO/ modENCODE) | Strain | Larval stage | Antibody | Temperature (°C) |
| --- | --- | --- | --- | --- | --- |
| **ChIP dataset** |  |  |  |  |  |
| HPL-2 ChIP-seq | GSE49749/modENCODE_5976 | N2 | L3 | HPL-2 SDQ2340 | 20 |
| **Histones datasets** |  |  |  |  |  |
| H3K9me1 | GSE25353 | N2 | L3 (mixed Male and Hermaphrodite population) | AB9045 | 20 |
| H3K9me2 | GSE49728 | N2 | L3 (mixed Male and Hermaphrodite population) | HK00008 | 20 |
| H3K9me3 | GSE49719 | N2 | L3 (mixed Male and Hermaphrodite population) | AB8898 | 20 |
| H3K27me3 | GSE49724 | N2 | L3 (mixed Male and Hermaphrodite population) | UP07449 | 20 |
| BLMP-1 | GSE25803/modENCODE 2612 | OP109 | L1 | GFP::3xFlag | 20 |

